# Variant-specific nucleocapsid mutations shape host innate immune trajectories during SARS-CoV-2 infection

**DOI:** 10.64898/2026.09.03.749134

**Authors:** Julia M. Berger, Jonas Schröder, Timothy K. Soh, Neele Pekarek, Lara- Marie Breul, Simon Dörner, Patrick Blümke, Jiabin Huang, Jens B. Bosse, Susanne Pfefferle, Charlotte Uetrecht

**Author notes:** corresponding authors (C.U.), (S.P.), (J.B.B.). these authors contributed equally (Julia M. Berger, Jonas Schröder). these authors contributed equally (Jens B. Bosse, Susanne Pfefferle, Charlotte Uetrecht).

## Abstract

The SARS-CoV-2 nucleocapsid (N) protein is one of the most highly expressed proteins during infection and plays crucial roles in the protection and packaging of viral RNA, replication, suppression of the immune response and virus assembly. The N gene and the overlapping accessory open reading frame ORF9b have continually evolved throughout the circulation of the virus, likely due to ongoing adaptation to the human host. This genetic variability influences the interplay of the N protein and its interactors. Yet, little is known about how specific mutations within the N/ORF9b locus of Variants of Concern (VOCs) shape the progression and outcome of infection. Here, we use a multi-omics approach to decipher how these genetic alterations reprogram the host cell by creating Wuhan-Hu-1-based recombinant viruses carrying an isogenic backbone with respective N mutations from the VOCs (called rNs) and comparing their effects at the transcriptome and proteome levels. We found that the mutations induce distinct transcriptional and translational alterations: rN-Alpha drives a stealth-like infection characterized by sustained translation efficiency and specific evasion of the 2’-5’-oligoadenylate synthetase (OAS) innate immune sensor, whereas rN-Delta and rN-BA.2 trigger a highly elevated inflammatory response. For rN-Delta, hyperphosphorylation of the N-protein drives cellular stress culminating in necroptotic cell death. Because these viruses differ only within the N/ORF9b locus, N sequence variation emerges as a determinant of infection outcome in its own right, warranting increased surveillance attention.

## 1. Introduction

The severe acute respiratory syndrome coronavirus 2 (SARS-CoV-2) nucleocapsid (N) protein is among the most abundant viral proteins produced during infection and fulfills essential roles across the viral life cycle. Beyond its canonical function in ribonucleoprotein complex assembly and genome packaging, N participates in viral RNA replication, modulates liquid-liquid phase separation (LLPS) to form replication organelles^1–4^, and suppresses host stress granule formation by interaction with host protein Ras GTPase-activating protein-binding protein 1 (G3BP1)^5–7^. N also engages multiple host signaling cascades: it inhibits interferon-β (IFN-β) production, activates NF-κB, and triggers NLRP3 inflammasome signaling, establishing it as a central node in both viral replication and innate immune evasion^8,9^. This multifunctionality positions N as a key determinant of the host-virus interaction landscape.

Genomic surveillance has shown that mutations within the viral genome concentrate predominantly in the spike, the non-structural protein (nsp) 12 and the nucleocapsid genes, with N being the third most commonly mutated protein of the virus^10,11^. Over the course of the pandemic, mutations accumulated within the gene, ultimately settling into a mutational profile characteristic of the Omicron lineages^12^. Several individual N mutations have already been linked to measurable phenotypic consequences, such as the R203K/G204R double substitution^13^ and the G215C substitution in the Delta variant of concern (VOC)^14^, indicating that even single amino acid changes in N can have functional effects. Importantly, the N gene partially overlaps with the open reading frame (ORF) 9b, a known innate immune antagonist^15,16^, meaning that mutations in N can simultaneously alter the ORF9b protein product^17^.

Despite a growing appreciation of the evolutionary trajectory of N, a thorough understanding of how VOC-specific N mutations reprogram the host cell has not yet been reached. A major limitation is that most available functional data on N are derived from overexpression systems, which lack the stoichiometric and temporal context of a genuine infection^18–28^. While interactome and multi-omics studies have mapped hundreds of interactions and demonstrated that VOC evolution shapes innate immune evasion^21,29^, attributing these system-level effects specifically to N sequence variation rather than to the full constellation of variant mutations has proven difficult.

To isolate and test the specific contribution of N mutations within a controlled viral backbone, we here established a matched reverse-genetics panel of recombinant viruses based on the ancestral Wuhan-Hu-1 backbone, in which the wild-type N open reading frame (ORF) was systematically exchanged with the corresponding sequences from the Alpha (B.1.1.7), Delta (B.1.617.2), BA.2 (B.1.1.529.2) and Pirola (BA.2.86) VOCs. By combining transcriptomics, proteomics and immunoprecipitation-mass spectrometry (IP-MS) of infected human lung epithelial cells, we identified differentially regulated genes, proteins, and interaction partners driven exclusively by the N/ORF9b cassette.

This design links variant-specific N/ORF9b sequences directly to host-state changes, decoupling specific functional effects of the N protein from those of the overlapping ORF9b, and reveals connections to differential antiviral and inflammatory trajectories, antigen-presenting behavior, and broader metabolic networks.

## 2. Materials and Methods

### 2.1 Cells

Vero E6, A549-ACE2-TMPRSS2 and Calu-3 cells were cultured under standard cell culture conditions as described before^30^.

### 2.2 Reverse genetics

#### Expression Plasmids

Viral RNA of wild-type SARS-CoV-2 from patient isolates was extracted with the QIAamp Viral RNA Kit for RNA extraction (QIAGEN, 52904). The N gene was then converted to cDNA and amplified with the SuperScript II One Step RT-PCR system (Thermo Fisher Scientific, 10928- 034). The linker of the pCC1-4k-Wuhan-Hu-1 nano BAC, including the 3’ end of the viral sequence starting at ORF10, was amplified with Q5 polymerase. Fragments were assembled with the NEBuilder (NEB, E2621S) according to manufacturer’s protocol. For primers see Supplementary Table 1. Assemblies were then transformed into Transformax Epi300 electrocompetent *E. coli* (LGC, 300110) and colonies were screened for N and the correctness of the N gene with the N insert fwd (5’CATGACGTTCGTGTTGTTTTAGA) and N insert rev (5’GCGAAAACGTTTATATAGCCCATCT) primers in a Q5 polymerase reaction. Expression plasmids were then isolated with the NucleoSpin Plasmid Kit (Macherey-Nagel, 740588.50). The N sequence and the presence of the mutations were confirmed with Sanger Sequencing (Microsynth).

#### Viruses

Rescue of the recombinant viruses was done as described previously^31,32^. Fragment amplification was done either off the pCC1-4k-Wuhan-Hu-1 nanoluc BAC, or off a previously created cDNA clone based on the pCC1-4k-Wuhan-Hu-1 with depletion of the BsaI sites^30^. For fragment amplification, Q5 polymerase (NEB, M0491S) was used according to the manufacturer’s protocol. The fragments containing the N gene, and its variant specific mutations, were amplified from the expression plasmids described above. For primers, see Supplementary Table 2. All fragments were sequenced to ensure correctness and presence of the VOC mutations in the N segment. Fragments were pooled at equimolar concentration of 0.1 pmol per fragment and were transfected with Lipofectamine 2000 (Invitrogen, 10696153) into HEK-293-ACE2 cells and then passaged onto Vero E6 cells for virus rescue.

### 2.3 Replication Kinetics

Cells were seeded and infected in triplicate with the recombinant viruses. Samples for titration were taken every 24 h, starting at time point 0 hpi until time point 72 hpi. Samples were then titrated on Vero E6 cells and graphed using Python.

### 2.4 RNA sequencing

Calu-3 cells were seeded in 6-well plates three days prior to infection. At confluency, they were infected with the recombinant viruses in triplicate at MOI 1. At 48 hpi, samples were inactivated in TRIzol reagent and RNA was extracted according to manufacturer’s protocol. High RNA integrity was confirmed via a TapeStation RNA ScreenTape analysis (Agilent, 5067-5576, 5067- 5577, 5067-5578). Per sample, 1 µg total RNA (quantified by Qubit RNA HS Assay, Thermo Fisher, Q32852) was Poly(A)-captured using the Lexogen Poly(A) RNA Selection Kit and further processed via the RNA-Seq V2 Library Prep Kit with UDIs (Lexogen, catalog number: 181.96) in the short insert size variant (RTM) according to manufacturer’s instructions (applying 11 cycles of Library Amplification PCR, step 4.3, User Guide version 171UG394V0111). Libraries were quality controlled on a TapeStation D5000 Assay (Agilent, D5000 ScreenTape 5067-5588 with D5000 Reagents 5067-5589) and were sequenced on an Element Biosciences AVITI instrument (2x75 Sequencing Kit Cloudbreak Freestyle Medium Output, product number: 860-00014) in a paired-end mode (2 x 80 bp). Following demultiplexing, each sample yielded 25.5-40.6 million raw read pairs. Adapter removal and quality filtering were performed using Trimmomatic (v0.36) to clip adapters, trim low-quality bases from read ends, apply sliding-window quality trimming (4-bp window, average Q ≥ 15), and discard reads shorter than 36 bp. Downstream processing was executed using the nf-core/rnaseq pipeline (v3.12.0)^33^ orchestrated by Nextflow (v22.10.5)^34^. Reads were aligned to the GRCh38 reference genome using STAR (v2.6.1d)^35^, followed by transcript-level quantification with Salmon (v1.10.1)^36^ using alignment-derived splice junction information. Transcript-level abundances from Salmon were aggregated to gene-level counts using tximeta (v1.12.0)^37^. Raw counts were normalized and variance-stabilized using DESeq2’s regularized log (rlog) transformation. To assess global transcriptomic structure, principal component analysis (PCA) was performed on the 500 most variable genes, confirming clear separation by variant identity and high reproducibility across biological replicates. Differential expression analysis was conducted using DESeq2 (v1.28.0)^38^. Genes were classified as significantly differentially expressed (DEGs) when they satisfied thresholds of |log₂ fold change| ≥ 2 and a Benjamini-Hochberg adjusted p-value ≤ 0.05. The rlog-transformed expression values were subsequently used for hierarchical clustering and downstream visualization.

### 2.5 Bottom-up sample preparation of infected lysates

Calu-3 cells were cultured in T75 flasks and infected at an MOI of 1. After 48 h of incubation, cells were washed twice with ice-cold phosphate-buffered saline (PBS) to remove extracellular contaminants. Cell pellets were subsequently lysed in 1 mL of lysis buffer supplemented with 1% Triton X-100. For protein recovery, a 100 µL aliquot of the lysate was combined with 400 µL of ice-cold acetone and incubated overnight at -20 °C. The resulting precipitate was collected by centrifugation at 21,300 × *g* for 1.5 h. The supernatant was discarded, and the protein pellets were air-dried at room temperature. Dried pellets were resuspended in 30 µL of a denaturation buffer consisting of 8 M urea in 100 mM Tris-HCl. The protein solution was diluted by the addition of 45 µL of 100 mM Tris-HCl. Disulfide bonds were reduced by adding 1 µL of 0.5 M DTT and incubating at 37 °C for 30 min. Subsequent alkylation was performed by adding 5 µL of 0.2 M iodoacetamide (IAA), followed by a 40-min incubation at room temperature in the dark. For enzymatic digestion, the samples were further diluted with 130 µL of 100 mM Tris-HCl to reduce the urea concentration, followed by the addition of 2 µL of trypsin. Digestion was carried out overnight at 37 °C under constant agitation at 600 rpm. The reaction was quenched and peptides were stabilized by the addition of 50 µL of stage-tipping buffer (10% acetonitrile (ACN) and 3% trifluoroacetic acid (TFA) in deionized water (18 MΩ·cm; Merck Millipore). The final peptide extracts were stored at -80 °C prior to liquid chromatography-tandem mass spectrometry (LC-MS/MS) analysis.

### 2.6 Immunoprecipitation

Immunoprecipitation of the viral nucleocapsid protein was performed using Dynabeads™ Protein G (Cat. No. 10004D, Invitrogen). Prior to IP, 1250 µL of Dynabeads were conjugated with a rabbit polyclonal anti-nucleocapsid antibody (Invitrogen, Cat. No. PA5-116894, Lot AC4660029) by incubating the beads in 5 mL of D-PBS-T (0.1%, Gibco) containing 0.1% bovine serum albumin (BSA, Roth Cat No. 2834.3) and 3 µg of antibody per 100 µL. This mixture was incubated for 30 min at room temperature under continuous rotation. For each sample, 28 µL of the antibody-conjugated beads were aliquoted and washed once with 200 µL of D-PBS containing 0.1 % BSA. Subsequently, the remaining 400 µL of the cleared cell lysate was added to the prepared beads and incubated for 2 h at 4 °C with rotation. Following incubation, the beads were washed three times with PBS. To minimize background contamination, the beads were transferred into clean reaction tubes prior to elution. Bound proteins were eluted from the beads by adding 30 µL of an elution buffer consisting of 8 M urea in 100 mM Tris. The elution was carried out for 25 min at room temperature. All samples (prepared in three replicates per condition for downstream bottom-up MS) were immediately frozen and stored at -80 °C until further analysis.

### 2.7 LC-MS/MS Analysis

Peptides were desalted and concentrated using EvoTips (Evosep, Odense, Denmark) according to the manufacturer’s instructions. Briefly, tips were conditioned and acidified, followed by the loading of 5 µL of sample in 15 µL of Buffer A (0.1% formic acid in H_2_O). Chromatographic separation was performed on an Evosep One system using the standardized SPD30 (30 samples per day) method with a 44-min gradient. Peptides were separated on a 15 cm Endurance Column (EV1113).

The Evosep One was online-coupled to a timsTOF Pro mass spectrometer (Bruker Daltonics, Bremen, Germany) equipped with a CaptiveSpray ion source. The instrument was operated in Data-Dependent Acquisition (DDA) mode using Parallel Accumulation-Serial Fragmentation (PASEF) technology or Data-independent acquisition (DIA) mode for full proteome samples.

### 2.8 Data analysis

#### Protein Identification and quantification

Raw DIA MS files were processed using DIA-NN software 2.0 Academia^39^. Prior to empirical data extraction, an *in silico* spectral library was generated using DIA-NN’s integrated deep learning-based spectra and retention time prediction engine. This prediction was based on a combined FASTA database comprising the human reference proteome (UniProtKB, canonical isoforms, downloaded October 2019) concatenated with the SARS-CoV-2 reference proteome (isolate Wuhan-Hu-1), including non-structural proteins (nsps) and specific variant sequences. For the library generation, trypsin was defined as the proteolytic enzyme with a maximum of one missed cleavage. The theoretical search space was constrained to peptide lengths of 7 to 30 amino acids, precursor charge states of 2 to 4, a precursor mass-to-charge (*m*/*z*) range of 350 to 1150, and a fragment *m*/*z* range of 200 to 1800. Cysteine carbamidomethylation was set as a fixed modification, while methionine oxidation and phosphorylation of serine, threonine, and tyrosine (STY) residues were configured as variable modifications (restricted to a maximum of one variable modification per peptide).

For the subsequent targeted extraction of the empirical DIA data, search parameters were adapted to encompass precursor charge states from 1 to 4, a precursor *m*/*z* range of 300 to 1800, and a fragment *m*/*z* range of 200 to 1800. Variable modifications for the main search included methionine oxidation, protein N-terminal acetylation, and N-terminal methionine excision. To ensure robust and consistent quantification across the entire dataset, the cross-run analysis (Match Between Runs) and “Smart Profiling” algorithms were enabled. The false discovery rate (FDR) was strictly controlled at 1% (q-value < 0.01) at both the precursor and protein group levels. Final protein quantification matrices were generated directly by DIA-NN, stringently filtered for 1% global and run-specific FDRs.

Raw DDA files were processed using the FragPipe (v24.0) computational platform^40^ (University of Michigan). Search tasks were executed against a concatenated database consisting of the human proteome (UniProtKB/Swiss-Prot, Taxonomy ID: 9606) and the SARS-CoV-2 proteome, supplemented with common laboratory contaminants.

Peptide identification was performed using the MSFragger (v4.4.1) search engine^41^. Digestion was set to strict trypsin with a maximum of two missed cleavages, restricting peptide lengths to 7–50 amino acids. For label-free quantification (LFQ), IonQuant (v1.11.20) was used with standard settings to ensure robust cross-sample comparison^42^. The FDR was strictly controlled at < 1%.

#### Statistical analysis and visualization

Protein intensities were log_2_-transformed, with missing values omitted. To balance robust quantification, proteins were filtered to require ≥2 valid observations in at least one experimental condition. To estimate condition-specific log-fold changes (LFC) while correcting for sample-level variance, a Bayesian hierarchical model was implemented in PyMC^43^ (v. 5.28.5). Observed intensities were modeled using a robust Student-t likelihood to minimize the influence of analytical outliers. To regularize LFC estimates across the high-dimensional dataset, a non-centered Horseshoe shrinkage prior was applied to the condition effects, effectively shrinking noisy, near-zero effects while preserving true, large shifts. Inference was performed using the No-U-Turn Sampler (NUTS) accelerated via the JAX/NumPy^44^ (v0.7.2/v2.0.2) backend. Posterior distributions of the condition-specific LFCs were summarized using the posterior mean and the 95% High-Density Interval (HDI) calculated via ArviZ^45^ (v0.22.0). Further data manipulation, statistical analyses, and high-dimensional visualizations including hierarchical heatmaps and multi-omics correlation plots were performed using custom in-house Python scripts. These computational workflows utilized the Pandas (v2.1.4, The pandas development team 2026) and NumPy libraries for data processing, SciPy^46^ and Scikit-learn^47^ for clustering and statistical metrics, and GSEApy^48^ for automated functional enrichment analysis. Graphical representations were generated using Matplotlib^49^ (v3.10.7) and Seaborn^50^ (v0.13.2).

#### IP-MS protein-protein interaction analysis

To account for technical variability in immunoprecipitation (IP) efficiency across different samples, the MaxLFQ intensities of identified proteins were normalized to the abundance of the bait protein (N). Specifically, log_2_-transformed MaxLFQ intensities of the N peptides were extracted, and the mean log_2_ intensity was calculated for each sample. Sample-specific correction factors (offsets) were determined by calculating the difference between the global mean of the bait across all IPs and the individual sample mean. These offsets were subsequently added to the log_2_-transformed intensities of all detected prey proteins in the respective samples. PBS negative controls were excluded from bait-normalization to preserve the true background signal.

Statistical analysis. Downstream data processing and statistical evaluation were performed using Perseus software (v2.1.4.0)^51^. Replicate reproducibility and overall sample clustering were evaluated using Pearson correlation coefficients. Missing values were imputed from a normal distribution simulating the limit of detection (width 0.3, down shift 1.8) to allow for statistical analysis of low-abundant or condition-specific interactors.

To identify significantly enriched interaction partners compared to the controls, two-sample Student’s t-tests were performed. Significance thresholds were established using a permutation-based FDR calculated with 100 randomizations. Interactors were stratified into two confidence tiers: High-confidence hits (Class A) were defined by a stringent FDR of 1% and an artificial variance parameter (S0) of 0.1 to penalize small fold changes. Medium-confidence hits (Class B) were defined by a relaxed FDR of 5%.

## 3. Results

### 3.1. Generation and replication kinetics of N/ORF9b-variant recombinant viruses

To isolate the contribution of the N/ORF9b sequence evolution to viral phenotype and host response, we used reverse genetics approaches^31,32^ to generate recombinant SARS-CoV-2 (rN) based on a cDNA clone^52^ (Figure 1A). Recombinant viruses share the ancestral Wuhan-Hu-1 backbone, as well as a nano-luciferase reporter in the ORF7ab. In additionally to the ancestral N/ORF9b sequence, recombinant viruses were created where the N/ORF9b sequence was replaced with the corresponding sequence of a wild-type isolate of the following VOCs: B.1.1.7 (Alpha), B.1.617.2 (Delta), B.1.1.529.2 (BA.2) and BA.2.86 (Pirola). Thus, the recombinants contain an identical genetic background, except for the specific mutations found in the N/ORF9b gene. In total, we rescued five recombinant viruses, namely rN-Wuhan, rN-Alpha, rN-Delta, rN- BA.2 and rN-Pirola. As the N and the ORF9b partially overlap in sequence, some mutations found towards the 5’ end of the N gene alter the ORF9b sequence; notably the amino acid sequence changes P13L, deletion (DEL)31/33, and D63G of N cause corresponding P10S, DEL27/29 and T60A mutations in the ORF9b (Table 1, Figure 1B).

**Figure 1.**
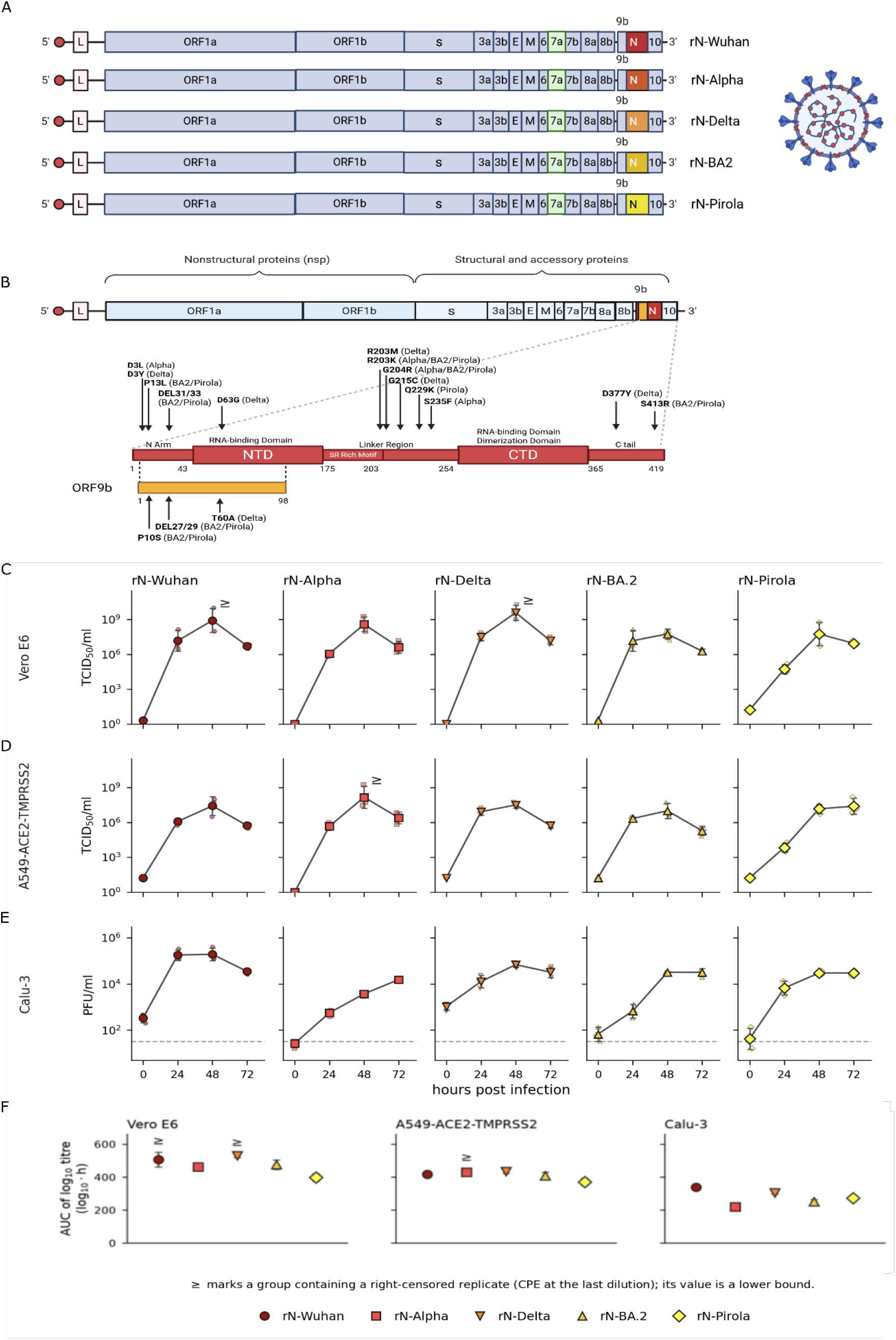
rN-VOCs and replication kinetics.. A - **Library of rescued recombinant viruses.** Recombinant viruses contain the Wuhan-Hu-1 backbone, containing a nano-luciferase reporter at the ORF7ab, as well as the N mutations found in the VOCs. In total, five recombinant viruses were rescued, including the rN-Wuhan, rN- Alpha, rN-Delta, rN-BA.2 and rN-Pirola. B - **Location and mutations of N and ORF9b within the SARS-CoV-2 genome.** N and ORF9b are located towards the 3’ terminus of the viral genome. Mutations found in the VOCs are indicated in bold, with corresponding VOC in which they occur in brackets. Mutations altering the ORF9b sequence are indicated in the same manner within the ORF9b. C-E - **Replication kinetics of recombinant viruses**. Vero E6 (c), A549-ACE2-TMPRSS2 (d) and Calu-3 (e) cells were infected in triplicates over a time course of 72 h for titration of viral titer. F - **Replication efficiency of recombinant viruses**. Replication efficiency of the individual recombinant viruses between the three cell lines are shown here. Significance values are given as AUC calculated through an unpaired t-test.

**Table 1.** Amino Acid and Nucleotide Changes affecting the ORF9b in comparison to the Wuhan- Hu-1 sequence.

| Amino Acid Change in N | Amino Acid Change in ORF9b | Nucleotide Change in N | Nucleotide Change in ORF9b | Affected Variant of Concern |
| --- | --- | --- | --- | --- |
| P13L | P10S | CCC→CTC | CCC→TCC | Omicron (BA.2/Pirola) |
| DEL31/33 | DEL27/29 | GAACGCAGT→<br>/ | AACGCAGTG→<br>/ | Omicron (BA.2/Pirola) |
| D63G | T60A | GAC→GGC | ACC→GCC | Delta |

To assess the replicative fitness of the N/ORF9b recombinants, we followed virus production over 72 h in interferon-deficient Vero E6 cells and in the interferon-competent cell lines A549- ACE2/TMPRSS2 and Calu-3, quantifying infectious progeny by endpoint titration (Vero E6 and A549-A/T) and plaque assay (Calu-3) (Figure 1C-E).

All five recombinants replicated productively in all three cell lines. Infectious titers increased markedly during the first 24–48 h and generally reached their maximum or a plateau by 48–72 h post infection. Thus, exchange of the N/ORF9b region did not abolish replication in any of the recombinants tested.

The recombinants nevertheless differed substantially in the magnitude and kinetics of infectious virus production. When individual growth curves were summarized as the area under the log₁₀ titer curve (Figure 1F), AUCs were higher in Vero E6 than in A549-A/T cells for all five recombinants. This difference reached statistical significance only for rN-Delta (mean AUC 529 versus 433 log₁₀·h; Welch’s t-test, p = 0.0004; Holm-adjusted p = 0.002; n = 3). In Calu-3 cells, rN-Wuhan showed the highest overall infectious virus production and reached an early plateau by 24 h, whereas the variant N/ORF9b recombinants generally accumulated infectious progeny more gradually, with maximal titers reached at 48–72 h. Importantly, however, the relative replication phenotypes differed between cell lines, with no single recombinant consistently displaying the highest or lowest replication across all three systems. Thus, rather than defining a uniform fitness hierarchy, exchange of the N/ORF9b region resulted in pronounced, cell-type-dependent replication phenotypes.

### 3.2. Lineage-specific N mutations uncouple ORF9b abundance and dictate global viral translation

To establish how variant-specific N/ORF9b mutations alter the transcription and translation dynamics of the virus itself, we evaluated both absolute viral RNA transcript levels and translated viral protein abundance in Calu-3 cells infected at an MOI of 1 at 48 hpi using transcriptomics (polyA mRNA-seq) and quantitative bottom-up proteomics via DIA coupled with Bayesian model analysis.

At the transcription layer, quantitative RNA sequencing revealed a broad trend of transcriptional attenuation in the rN-Alpha variant. While normalized read counts for genomic RNA and the N- gene were consistently lower than those of the other recombinants, this suppression was statistically significant only in the 5’ leader sequences (Figure 2A). To analyze template-switching mechanics, we calculated N-gene and 5’ leader sequence ratios relative to genomic RNA (gRNA). The N/gRNA ratio remained stable across all recombinants, indicating conserved baseline N- gene transcription. For the leader/gRNA ratio, the rN-Delta exhibited a noticeably lower mean, though high inter-replicate variance precluded statistical significance. This is in line with findings that the R203M mutation significantly increases viral particle assembly and RNA encapsidation^53^, likely restricting subgenomic template switching in favor of continuous genomic replication (Figure 2B).

**Figure 2.**
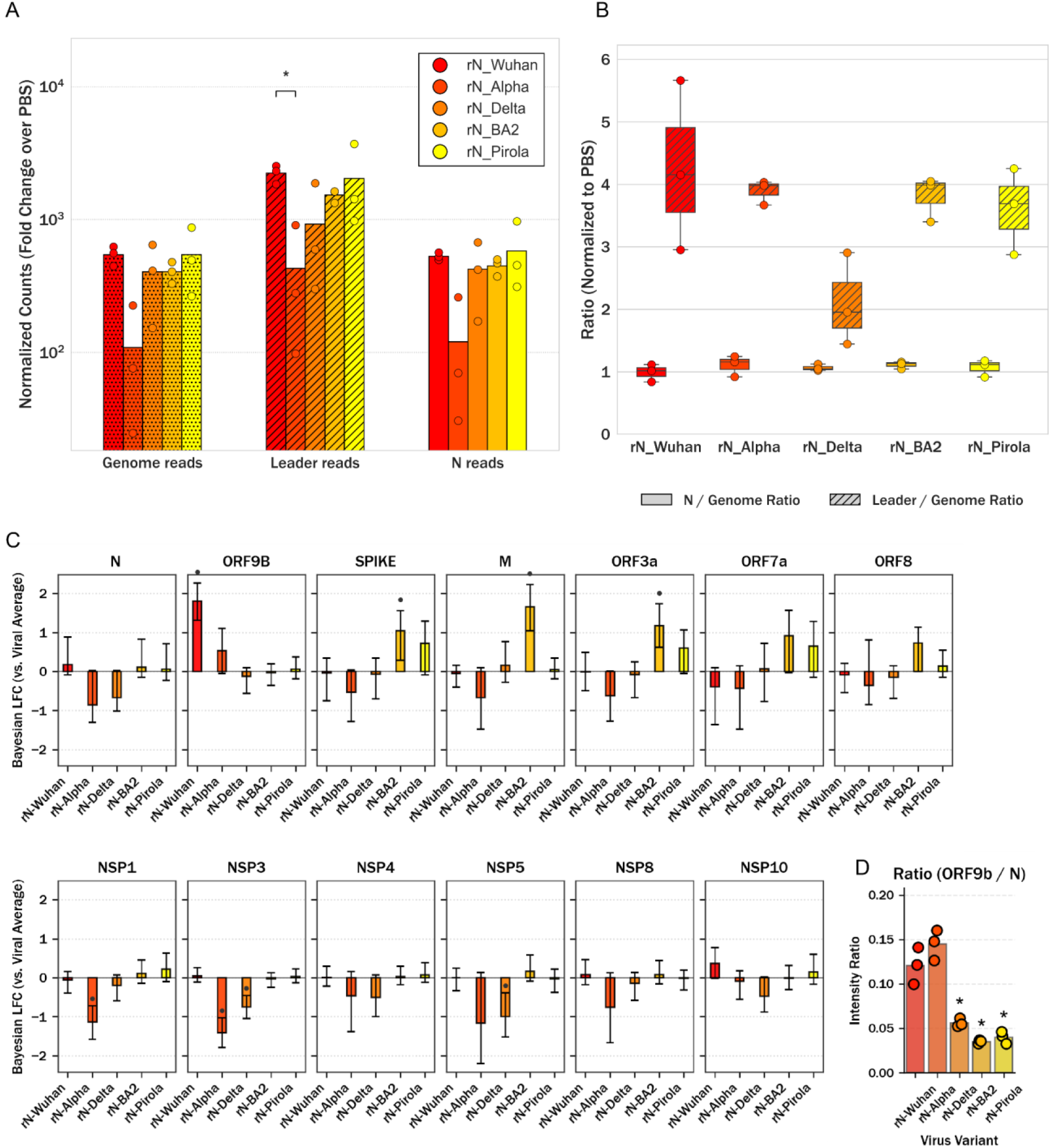
N transcript and protein abundance. A - **Viral RNA transcription and genomic load.** Quantification of viral RNA species across variants at 48 hpi (MOI 1). Values represent full genome reads, leader sequences, and N-gene transcripts, all normalized against the background PBS control to determine relative fold change or abundance. Data illustrates the efficiency of viral replication and subgenomic transcription across the different variants. Statistical significance was determined using multiple unpaired t- tests followed by a Bonferroni correction. B - **Viral RNA transcription and template-switching ratios.** Ratios of N-gene and 5’ leader sequence reads relative to full genomic RNA (gRNA) at 48 hpi (MOI 1), normalized to the PBS control. Boxplots display the median and interquartile range, with individual biological replicates (n=3) overlaid as scatter points. Statistical significance was evaluated on log_2_-transformed ratios using multiple unpaired t-tests followed by a Bonferroni correction C - **Condition-specific effects on viral protein expression.** LFC of quantified SARS-CoV-2 viral proteins compared to the global mean. Bar heights represent the posterior mean LFC, and error bars indicate the 95% HDI. D - **Depletion of steady-state ORF9b relative to N in VOCs.** MaxLFQ intensity ratios of ORF9b to N across rN-VOCs (Calu-3 cells, 48 hpi). Bars represent the mean of three biological replicates (scatter points). Statistical significance vs. rN-Wuhan was assessed via Welch’s t-test (* p<0.05).

We observed an extensive transcriptomic attenuation in rN-Alpha, which correlated directly with its viral translation profile (Figure 2C). While the N abundance itself was slightly lower in rN-Alpha and rN-Delta, this decrease was not statistically significant. However, the rN-Alpha recombinant exhibited a systemic, significant down-regulation of the rest of the viral replication machinery. It showed strong negative Bayesian LFC for the host-translation inhibitor nsp1 (LFC = -1.138), the papain-like protease and RNA pore nsp3 (−1.413), and the main protease nsp5 (−1.166). In contrast, the highly fit rN-BA.2 variant maintained robust, stable viral RNA transcript levels and showed an optimized translational shift.

Furthermore, we conducted quantitative ratio analysis that revealed a distinct stoichiometric regulation between the overlapping gene products N and ORF9b (Figure 2D). While the relative abundance ratio of ORF9b to N remained consistently high and virtually identical between the rN- Wuhan and rN-Alpha variants, we found a depletion of ORF9b relative to the viral proteome in rN-BA.2 and rN-Pirola. Because ORF9b occupies an alternative reading frame (+1) of the N gene and is therefore also affected by the DEL31/33, this may lead to structural alterations in the Omicron ORF9b core structure and negatively impact protein stability or half-life, potentially explaining the observed depletion. Due to our reverse genetics approach, these quantitative proteomics data allow us to see these direct biochemical consequences of the N/ORF9b mutations.

### 3.3 N-protein evolution drives divergent evasion of early innate immune sensing

To analyze how the individual rN-VOCs modulate global cellular homeostasis, we integrated the transcriptomic and proteomic landscape of infected Calu-3 cells. To map the variant-specific multi- omics landscape, we plotted the individual RNA and protein LFC. As shown in Figure 3A, rN- Delta and rN-BA.2 drive a concordant upregulation of classical innate immune factors like OAS1 and MX1. In contrast, the rN-Alpha maintains these genes significantly closer to baseline, reflecting a strong suppression of the host response. An initial visual assessment of this multi- omics scatter presents a methodological paradox regarding the rN-Alpha recombinant. While variants such as rN-Delta and rN-BA.2 exhibit broad, scattered profiles indicative of severe virus- induced cellular stress and RNA-protein uncoupling, the rN-Alpha displays a narrow, tightly clustered profile lacking extreme outliers. We analyzed the absolute LFC magnitudes to resolve this: the restricted spread observed during the rN-Alpha infection is a consequence of a dampening of the early host response. Because rN-Alpha efficiently blocks the initial transcriptional alarm, the downstream uncoupling of RNA and protein appears deceptively minimal (Supplementary Figure 1).

**Figure 3.**
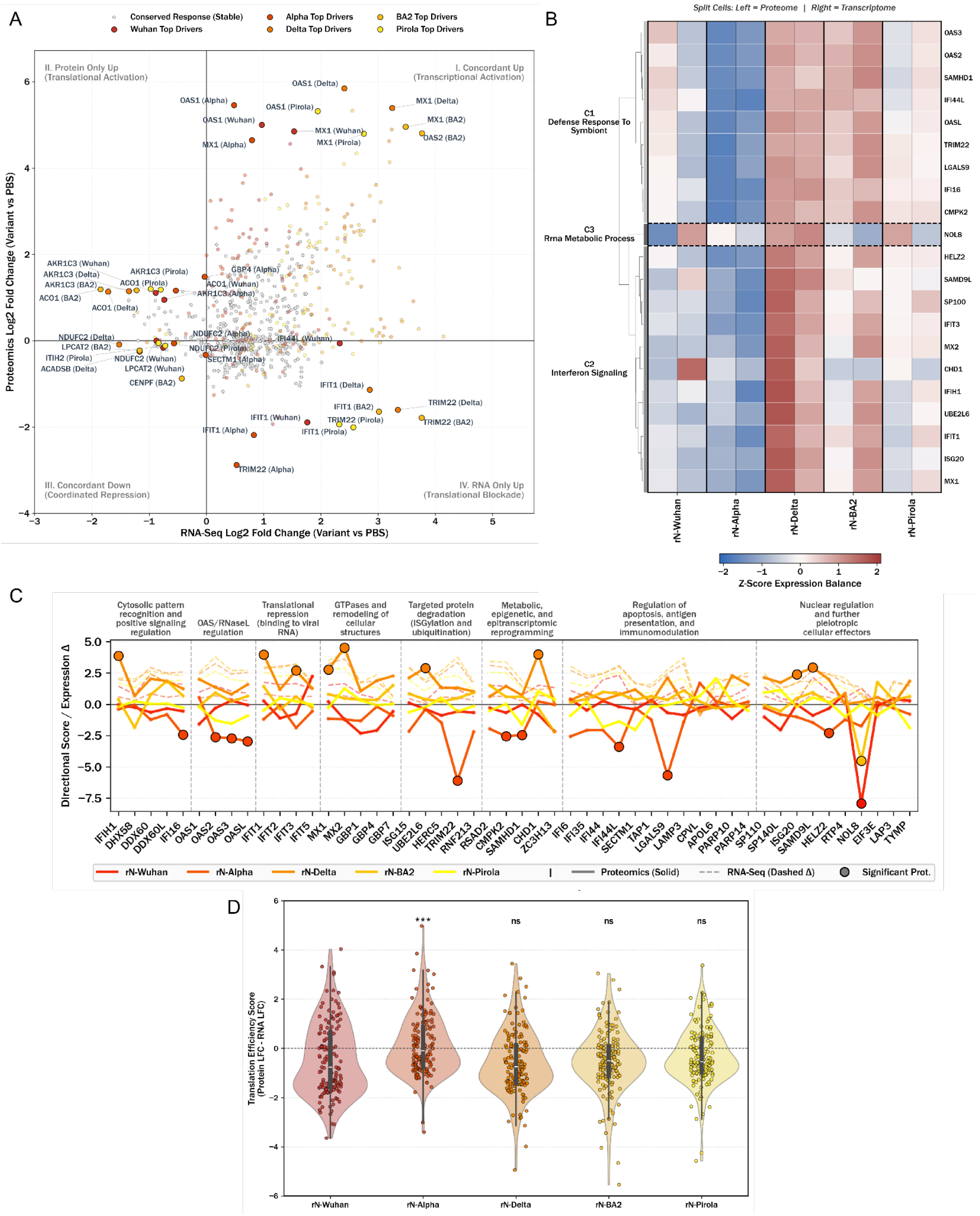
rN-VOC specific effects. A - **Multi-Omics landscape of host transcriptomic and proteomic responses.** Quadrant scatter plot correlating host transcriptomic (RNA-Seq, x-axis) and proteomic (y-axis) LFC relative to the mock (PBS) control for each SARS-CoV-2 recombinant variant at 48 hpi. Host factors exhibiting stable, conserved expression across all conditions (cross-variant variance ≤ 75^th^ percentile in both omics layers) are displayed as a grey background cloud. Variant-specific dynamic responses are colored according to the respective viral variant. I. Concordant Up (Transcriptional Activation), II. Protein Only Up (Translational Activation), III. Concordant Down (Coordinated Repression), and IV. RNA Only Up (Translational Blockade). The top two expression drivers per variant within each quadrant are highlighted and explicitly labeled. B - **Integrated Dual-Omics profiles and functional clustering.** Split-cell heatmap displaying matched host proteomic (left) and transcriptomic (right) profiles across viral variants. Values represent row Z-scores of PBS-subtracted median LFC, restricted to genes passing a proteomic Bayesian 95% HDI threshold. Hierarchical clustering (Euclidean distance, Ward’s linkage) of the proteomic layer was thresholded at 45% maximum distance. Clusters are annotated with their top-enriched pathway via Gene Set Enrichment Analysis (GSEA, adj. p<0.05). C - **Pathway-specific protein expression profiles.** Directional Scores (LFC / SD) of individual host proteins grouped by biological pathways. Scores are calculated by dividing the median LFC by the estimated standard deviation (derived from the 95% HDI). Large, black-rimmed markers denote statistically significant alterations (95% HDI excluding zero). Dashed lines indicate transitions between pathways. D - **Global RNA-protein expression divergence across viral variants.** Violin plot displaying the distribution of expression divergence, calculated as ΔLFC=LFC_protein_−LFC_RNA_ relative to the PBS control. Internal horizontal lines indicate quartiles, and the dashed line at zero represents perfect omics concordance. Statistical significance for directional shifts was evaluated per variant using a two-tailed Wilcoxon signed-rank test against a median of rN-Wuhan (ns: p≥0.05, *: p<0.05, **: p<0.01, ***: p<0.001).

We further corroborated this distinct regulatory dichotomy by mapping the global expression patterns. We applied Euclidean hierarchical clustering to both the host transcriptomic and proteomic datasets, revealing distinct, variant-specific co-expression modules within a dual-omics expression matrix (Figure 3B). At the transcriptomic level, canonical type I/III interferon (IFN) and interferon-stimulated gene (ISG) defense programs were strongly induced by the rN-Delta and rN-BA.2 variants, but actively suppressed in the rN-Alpha variant. This transcriptomic dichotomy was mirrored at the proteomic layer, demonstrating that the global stress and antiviral signatures heavily upregulated by rN-Delta and rN-BA.2 are maintained in a tightly controlled, attenuated state during rN-Alpha infection.

To dissect this variant-specific modulation beyond global metrics, we resolved the multi-omics dataset into specific functional pathways, focusing on immune factors and IFN signaling (Figure 3C). Tracking the standardized protein abundance of individual functional factors across the viral evolutionary timeline revealed a highly synchronized, non-linear evolutionary trajectory. Across all signaling pathways of the innate immune system, the Ns of the recombinant alpha virus and the subsequent Omicron sublineage Pirola appear to have induced a suppressed, less inflammatory state. Most notably, rN-Alpha effectively evaded initial viral sensing by strongly suppressing the production of frontline antiviral sensor proteins, particularly those of the OAS/RNase L system, at both transcriptional and translational levels.

To pinpoint whether this suppression is driven by a global host shutoff or a specific upstream sensing evasion, we calculated global translation efficiency scores (Figure 3D). Remarkably, rN- Alpha did not exhibit a global translational block; instead, it showed significantly higher translation efficiency than the ancestral rN-Wuhan strain. The other variants maintained a baseline efficiency similar to the wild type. This difference demonstrates that rN-Alpha maintains a fully competent translational machinery. Therefore, this indicates that the phenotype exhibited by rN-Alpha is not achieved through downstream translational repression, but by successfully preventing the initial activation of the host’s antiviral transcriptional program. Notably, rN-Wuhan elicited a more robust innate immune response despite exhibiting the highest absolute ORF9b levels, indicating that the rN-Alpha phenotype does not simply scale with ORF9b abundance. Together, these data suggest that lineage-specific functional properties of the N/ORF9b module, rather than global host shutoff or ORF9b expression levels alone, shape the distinct innate immune phenotype of rN-Alpha.

### 3.4 Interactome remodeling mediates immune stealth via enhanced G3BP1 sequestration

To identify the precise upstream physical protein-protein interactions driving these profound downstream phenotypic shifts in immune evasion and cell fate, we performed IP-MS of infected Calu-3 cells at 48 hpi and MOI 1 against N across the recombinant viruses. We used a polyclonal anti-N antibody to ensure variant-independent capture of N across multiple epitopes, preventing pull-down biases that could arise from lineage-specific mutations.

We found that established N interactors such as G3BP1^7,54,55^ and Y-box binding protein 1 (YBX1)^56^ were prominently enriched across the variants, validating our IP-MS approach. While absolute fold changes confirmed robust enrichment across conditions (Supplementary Figure 2), we identified variant-specific binding preferences from bait-normalized relative interaction affinities (Figure 4A). Notably, the relative affinity for YBX1, which promotes translation of viral structural proteins, was highest during rN-BA.2 infection. Conversely, the bait-normalized enrichment of G3BP1, the primary nucleator for cytoplasmic stress granule assembly, reached the highest levels of the recombinants during rN-Alpha and rN-Delta infections (Figure 4A). We hypothesize that by strongly sequestering G3BP1, rN-Alpha can dismantle these stress granules, preventing both the translational shutdown caused by the spatial segregation of viral RNA and the subsequent local accumulation of pathogen-associated molecular pattern (PAMP)-sensing proteins such as RIG-I. Additionally, this correlates with the suppression of the early transcriptional ISG alarm captured in our transcriptomic data.

**Figure 4.**
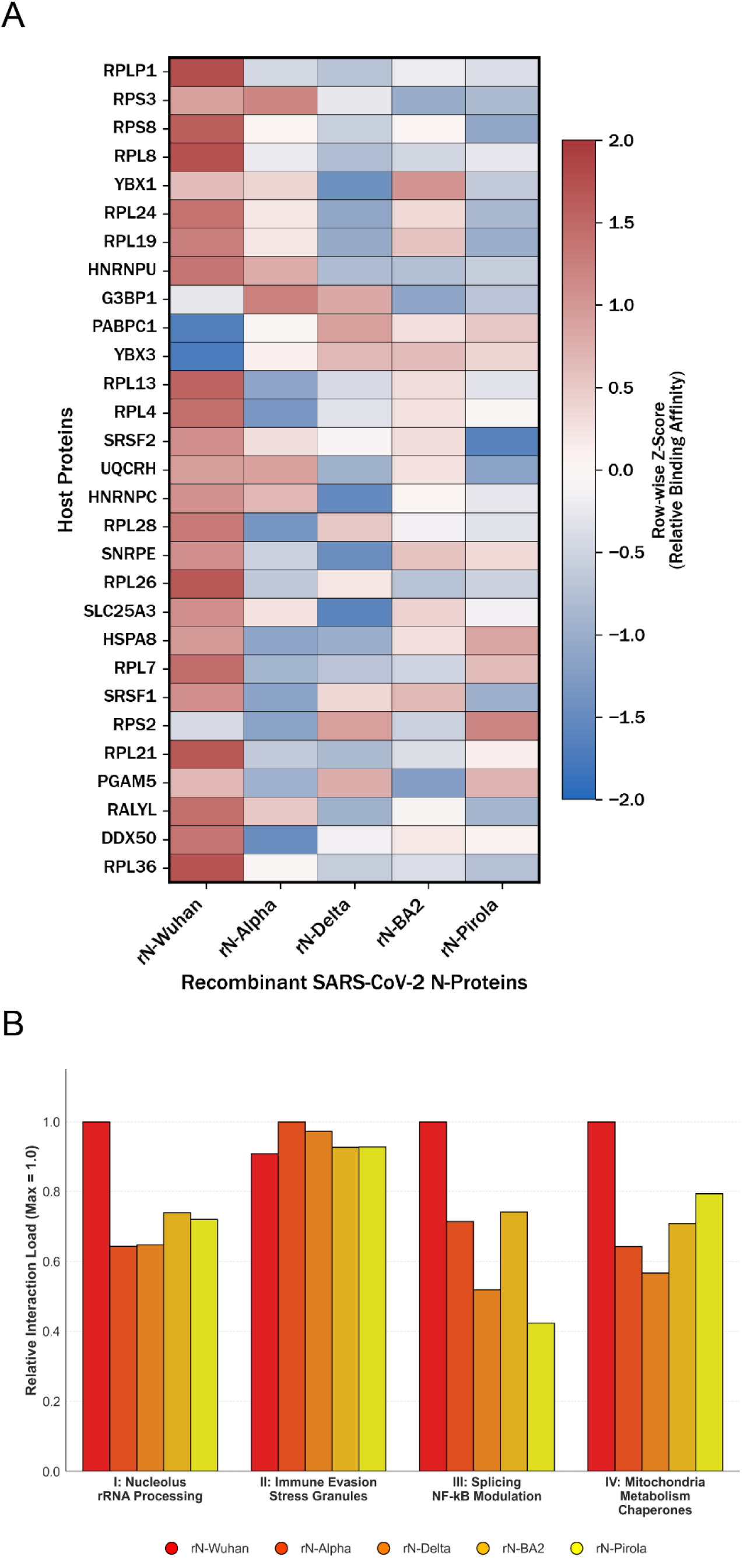
rN-VOC interactome. A - **Relative interaction affinities of high-confidence host proteins across recombinant SARS-CoV-2 N-protein variants**. Calu-3 cells were infected at an MOI of 1 and harvested at 48 hpi. Viral N complexes were isolated via IP-MS utilizing a polyclonal anti-N antibody. Values are displayed as row-wise Z-scores derived from bait-normalized LFC compared to the respective mock-infected (PBS) negative controls. Only host interactors exhibiting statistically significant enrichment (FDR < 0.05) in at least one recombinant condition are shown. B - **Relative host interactome load of SARS-CoV-2 N-protein recombinants.** Aggregated IP- MS LFC across functional clusters (I-V). Significant LFC values were summed per cluster and normalized to the respective maximum variant score (Max = 1.0), Supplementary Table 3.

Following this, we aggregated the N-protein interactors into functional clusters (Supplementary Table 3) and visualized the interaction fold changes (Figure 4B). This analysis corroborates the divergent evolutionary trajectories of the variants. The ancestral rN-Wuhan exhibited broad, high-affinity interactions with most of the host machinery, indicative of a highly disruptive host shutoff. In contrast, the ’stealth’ variant rN-Alpha displayed a minimal interactome footprint, reducing interactions with host ribosomes while maintaining peak engagement with the stress granule/immune evasion cluster. The rN-Pirola recombinant exhibited the lowest interaction with host splicing factors, supporting a host-preservation strategy, while re-engaging the host chaperone and mitochondrial networks. This elevated interaction may act as a critical buffer, enabling Pirola to manage translational burdens without triggering fatal endoplasmic reticulum (ER) stress.

### 3.5. N-terminal hyperphosphorylation switches host cell fate to inflammatory necroptosis

Given the distinct growth kinetics and immune evasion strategies among the variants, we evaluated whether these differences correlated with altered host cell fate. Proteomic quantification demonstrated that classical apoptosis was comparably induced across all variants, as we observed a uniform upregulation of executioner caspases (e.g., CASP3; Figure 5A) and intrinsic regulators like BAX, with no statistically significant differences between the recombinant viruses. This indicates that all Ns permit a baseline apoptotic response relative to uninfected cells.

**Figure 5.**
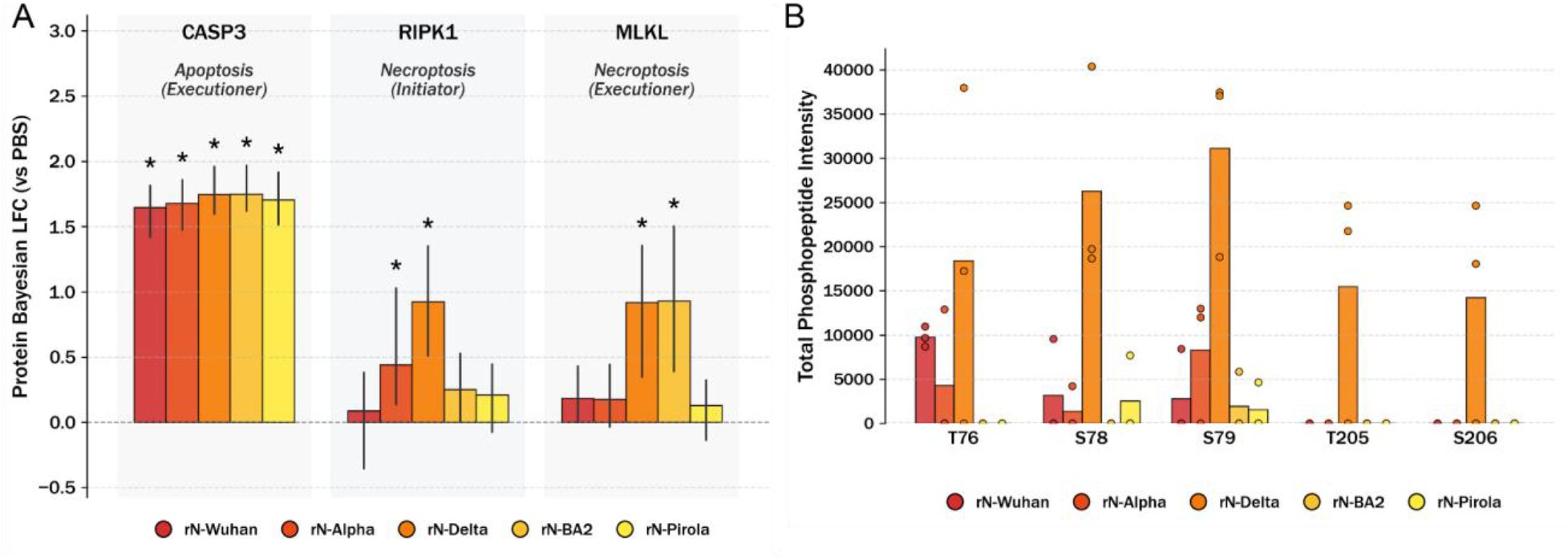
cell death pathways and correlation with N phosphorylation. A - **Cell death pathway divergence across viral variants.** Grouped bar chart displaying the Bayesian LFC of key apoptotic (CASP3) and necroptotic (RIPK1, MLKL) proteins relative to the PBS control. Error bars indicate the 95% HDI. Asterisks (*) denote statistically significant expression changes where the 95% HDI completely excludes the PBS baseline (normalized to zero). B - **Variant-specific N phosphorylation.** Bar graph displaying the total phosphopeptide intensity per specific residue across the recombinant viral variants. Intensities of detected phosphorylation sites were summed across all charge states to determine total abundance. Bars represent the mean intensity for each variant with individual biological replicates represented by overlaid scatter points.\

However, we found a statistically significant divergence in the regulation of inflammatory cell death. The rN-Delta recombinant triggered a robust necroptotic cascade, significantly upregulating the kinase RIPK1 and the terminal executioner MLKL (Figure 5A)^57,58^, driving the host cell toward chaotic, inflammatory lysis. The rN-Alpha, on the other hand, actively repressed this pathway, maintaining both RIPK1 and MLKL at significantly lower levels than rN-Delta.

To investigate the mechanisms driving this unique inflammatory response of rN-Delta, we analyzed the post-translational modification profiles of the rN-VOCs (Figure 5B). There, rN-Delta exhibited a higher proportion of phosphorylated N-peptides specifically within the highly conserved N-terminal region (residues T76, S78, and S79) compared to all other variants. This hyperphosphorylated state is likely a consequence of Delta-specific structural mutations, for instance G215C^14,59^, which stabilize dimer formation and alter the biophysical properties of the N-protein. These alterations elevate its propensity to form higher-order oligomeric LLPS assemblies. We hypothesize that these dense condensates then act as a recruitment platform for host kinases and their substrates, driving the observed hyperphosphorylation and cellular stress. Consequently, this stress overrides standard apoptotic signaling and triggers the observed necroptotic cascade. Furthermore, this necroptotic shift was independently reproduced in A549- ACE2-TMPRSS2 cells, confirming the robust nature of this variant-specific cell fate (**Supplementary Figure 3**).

This specific necroptotic divergence mirrors the non-linear evolutionary pattern observed in the innate immune response. The rN-BA.2 acts in a hybrid way, triggering the terminal necroptotic executioner MLKL as aggressively as rN-Delta. In contrast, the rN-Pirola recombinant restores the rN-Alpha-like evasion strategy by abrogating the necroptotic cascade. Collectively, these data suggest that successful viral adaptation epitomized by rN-Alpha and restored in rN-Pirola is achieved by specifically silencing necroptotic chaos to preserve cellular architecture for late-stage viral assembly.

## 4. Discussion

Our multi-omics analysis of the recombinant viruses demonstrates that evolution of the N/ORF9b locus is a major global regulator of viral replication dynamics and host immune evasion. We show that the VOC Ns pursue fundamentally distinct strategies: while rN-Alpha adopted a stealth-like phenotype characterized by marked suppression of the OAS response without global translational shutoff, rN-Delta induced a contrasting hyper-inflammatory state associated with necroptotic cell death. Within the Omicron lineage, we found a phenotypic recalibration: the early Omicron representative rN-BA.2 drives translation of structural proteins and hyper-inflammation at the expense of cellular stealth, whereas its successor the rN-Pirola effectively mitigates this pro-inflammatory phenotype through enhanced host preservation.

A critical advantage of our approach is the ability to functionally uncouple N from the known innate immune antagonist ORF9b^60,61^. The innate immune suppression observed in response to rN- Alpha infection is consistent with the enhanced silencing previously reported for the Alpha VOC^29^. The Alpha lineage features the D3L mutation and an upstream Kozak sequence alteration, which have been reported to optimize transcription regulatory sequence (TRS-B) and enhance leaky scanning to drive elevated ORF9b expression^29,55,62^. However, in our reverse genetics system, these upstream compensatory sequences are identical in the recombinant viruses, and the rN- Alpha was found to express lower levels of ORF9b protein than the rN-Wuhan does. Because rN- Wuhan triggers a more robust innate immune response despite possessing the highest absolute ORF9b levels, the profound immune silencing observed in rN-Alpha cannot be attributed to ORF9b abundance alone. Instead, our multi-omics comparison reveals that this ’stealth’ phenotype is driven by an upstream blockade orchestrated by the physical interactome of the Alpha N itself. By heavily sequestering G3BP1, Alpha effectively dismantles antiviral stress granules, preventing the subsequent local accumulation of PAMP-sensing proteins required to trigger the innate immune alarm, such as RIG-I^63^.

This direct, independent influence of N is further highlighted by the Omicron sublineages. The rN- BA.2 and rN-Pirola variants share the same ORF9b amino acid sequence, yet they exhibit contrasting phenotypes. The highly pronounced pro-inflammatory phenotype of rN-BA.2 aligns with the presence of the R203K/G204R double substitution, which increases replication, fitness, and pathogenesis, potentially through increased phosphorylation via proximal GSK-3 consensus sites^13^. The hyper-induction of the host’s innate immune system triggered by VOCs such as BA.2, and reflected by the rN-BA.2, can provoke a cytokine storm^64–67^ that damages the repair of lung epithelial cells^68^ and breaks down epithelial barriers, providing the virus with an optimal environment for transmission^69^. In contrast, rN-Pirola’s evasive recalibration mitigates this damage. A recent study has reported that Pirola’s Q229K substitution alters the host adaptive immune recognition^70^. Our data provide further insight into the role of Q229K, which drives effects on innate immune sensing, demonstrating that specific substitutions within N directly affect the broader host immune response.

The rN-Delta recombinant provides a counterpoint to the stealth strategy, illuminating how its replication triggers a strong cellular response. This finding aligns with published literature demonstrating that the hallmark Delta VOC mutation N:R203M significantly enhances viral assembly mechanics but heavily restricts template-switching kinetics in unadapted genetic backgrounds^53,71^. Interestingly, our dataset failed to reflect the anticipated inhibitory effect of rN- Delta on RIG-I-mediated interferon induction. Instead, rN-Delta drives a hyper-inflammatory response culminating in necroptosis, matching the higher disease severity observed in animal models compared to Omicron lineages and the Alpha lineage^72–74^. The literature presents conflicting reports regarding the post-translational modification patterns of the Delta N. While Syed et al.^75^ postulate that the R203M substitution attenuates phosphorylation, Yun et al.^76^ report no significant decrease, instead attributing hyperphosphorylation to the R203K/G204R mutations found in the Alpha and Omicron lineages. Our dataset provides a counter-narrative: the inflammatory escalation is intrinsically linked to the massive N-terminal hyperphosphorylation uniquely observed in rN-Delta. Together with the Delta-specific G215C mutation, which stabilizes dimer formation and alters biophysical properties to elevate RNA-binding and oligomerization^14,77^, this hyperphosphorylated state likely recruits host kinases into dense LLPS condensates^78,79^, driving profound cellular stress and subsequent necroptotic lysis.

Our study nevertheless has limitations. The replication kinetics revealed significant differences between cell lines of differing innate immune capacity, highlighting that host cell context shapes the response to the infection^67,74,80^. We therefore based the multi-omics comparison on Calu-3 cells, human lung epithelial cells that retain an intact interferon response, so that the host programs we describe are measured in a context permissive for innate immune signaling. Moreover, while we identified global interactomic shifts, the lineage-defining mutations (e.g., S235F, S413R, D377Y in N, and P10S, T60A in ORF9b) likely act through individual kinase-substrate relationships that our steady-state measurements do not resolve.

Taken together, our data establish the N/ORF9b locus as an independent determinant of how infection unfolds in the host cell. Exchanging this single locus in an otherwise identical genome moved the host response from a suppressed, low-inflammatory state to one that ends in necroptotic lysis, with no change to spike, the replicase or the remaining accessory proteins. Moreover, the resulting phenotypes were mostly dependent on which host proteins N engaged and its phosphorylation state rather than the amount of ORF9b produced. Importantly, the restraint of the inflammatory program has not accumulated steadily along the viral lineage: it is present in rN-Alpha, absent in rN-Delta and rN-BA.2, and re-established in rN-Pirola. N sequence variation is therefore a determinant of infection outcome in its own right, not simply carried along as variants evolve, and the surveillance and functional follow-up so far directed mostly at the spike protein should extend to this locus.

## 7. Acknowledgements

We thank Griffith University for providing the SARS-CoV-2 reverse genetics cDNA clone (received through https://mrcppu-covid.bio/). The following reagent was obtained through BEI Resources, NIAID, NIH: *Homo sapiens* embryonic kidney epithelial cells expressing transmembrane protease, serine 2 and human angiotensin-converting enzyme 2 (ACE2) (293T- ACE2.TMPRSS2 (mCherry), NR-55293).

## 8. Author Contributions

JMB - Conceptualization, Formal Analysis, Investigation, Methodology, Visualization, Writing - Original Draft Preparation

JS - Conceptualization, Formal Analysis, Investigation, Methodology, Visualization, Writing - Original Draft Preparation

TKS - Conceptualization, Formal Analysis, Investigation, Writing - Review and Editing NP - Investigation

PB - Investigation JH - Formal Analysis

JBB - Conceptualization, Funding Acquisition, Project Administration, Supervision, Writing - Review and Editing

SP - Conceptualization, Funding Acquisition, Project Administration, Supervision, Writing - Review and Editing

CU - Conceptualization, Funding Acquisition, Project Administration, Supervision, Writing - Review and Editing

## 9. Funding

The SP, CU, and JBB labs were funded by RTG 2771 Humans and Microbes project no. 453548970, and the CU and JBB labs were additionally funded by RTG 2887 VISION project number 497350882. The SP laboratory was also funded by CRC1328 (grant number 335447717). The JBB lab was also funded by the Deutsche Forschungsgemeinschaft (DFG, German Research Foundation) under Germany’s Excellence Strategy - EXC 2155 - project number 390874280, the CRC 1648 Emerging Viruses project number SFB 1648/1 2024– 512741711, the Research Unit FOR5200 DEEP-DV (443644894) project BO 4158/5-1 and BO 4158/5-2, the Research Unit FOR5898 AdBHealth (548065690) project BO 4158/9-1, and the German Center for Infection Research (DZIF) grants IICH TTU 07.918, IICH TTU 07.861 and IICH TTU 07.863 as well as the Wellcome Trust through a Collaborative Award (209250/Z/17/Z), Behörde für Wissenschaft, Forschung und Gleichstellung (BWFG) Hamburg through “Hamburg-X Infektionsforschung”, and the Leibniz ScienceCampus InterACt, funded by the BWFG Hamburg and the Leibniz Association (W75/2022). Equipment was funded by an equipment grant of the Behörde für Wissenschaft, Forschung, Gleichstellung und Bezirke (BWFG) of the Free and Hanseatic City of Hamburg to C.U. Instruments used in this study were funded by the BMFTR project VirMScan (BMFTR 13GW0622) and by a BMG Rapid Response grant.

## 10. Declaration of interests

The authors declare no competing interests.

## 11. Declaration of generative AI and AI-assisted technologies in the manuscript preparation process

During the preparation of this work, the authors used Opus5, Gemini 3.5 flash, GPT5.6 for language editing and literature research. After using these tools, the authors reviewed and edited the contents as needed and take full responsibility for the content of the published article.

## 12. Resource Availability

### Lead contact

Further information and requests for resources and reagents should be directed to and will be fulfilled by the lead contact, Jens B. Bosse.

## Materials availability

All recombinant viruses and plasmids generated in this study are available from the lead contact upon reasonable request.

## Data and code availability

The RNA sequencing data generated in this study have been deposited in the Gene Expression Omnibus (GEO) under the Series accession number GSE345656 (Sample accessions GSM10012437–GSM10012457; SRA accessions SRX35031754–SRX35031774).

The mass spectrometry proteomics data have been deposited to the ProteomeXchange Consortium via the PRIDE^81^ partner repository with the dataset identifier PXD083585 Any additional information required to reanalyze the data reported in this paper is available from the lead contact upon request.

## Notes

### Competing Interest Statement

The authors have declared no competing interest.

