## Supplementary materials for "Variant-specific nucleocapsid mutations shape host innate immune trajectories during SARS-CoV-2 infection"

### 6. Supplementary Material

Supplementary Table 1. Primers to create expression plasmids for reverse genetics fragments

| Primer Name | Primer Sequence | Fragment Length (bp) |
| --- | --- | --- |
| N BAC link fwd | tcccaggtaacaaaccaaccgatgtctgataatggacccc caaaatca | 1597 |
| SARS-CoV-2 CPER F6 rev | GTCATTCTCCTAAGAAGCTATTAAAATC ACATGGGG | |
| CPER BAC link fwd | TAGCTTCTTAGGAGAATGACAAAAAAAA AAAAAAAAAAAAA AAAAAAAAAGGGTCGGCATGGC | 5470 |
| CPER BAC link rev | GGTTGGTTTGTTACCTGGGAAGGTATAA ACCTTTAATCGGTT CACTAAACGAGCTCTGCT | |

Supplementary Table 2. Primers for fragment creation for recombinant virus rescue (Fig 2)

| | Primer Name | Primer Sequence | Fragment Length (bp) | Overlap length with next fragment (bp) |
| --- | --- | --- | --- | --- |
| rN-Wuhan rN-Alpha | GG1F | ggctacggtctcgTCCCAGGTAACAAACCAA CCAACTTTCG | 2874 | / |
| | GG1R | ggctacggtctccccacaacacaggcgaactc | | / |
| | GG2F | ggctacggtctccgtggcagatgctgtcataaaaac | 3599 | / |
| | GG2R | ggctacggtctcctctcctacaacttcggtag | | / |
| | GG3F | ggctacggtctccgagacattatacttaaaccagc | 3592 | / |
| | GG3R | ggctacggtctccgccatttttctaaaaccac | | / |
| | GG4F | ggctacggtctcctggcattcccatctggtaaag | 3581 | / |
| | GG4R | ggctacggtctccaaagtaagaatcaattaaattgtcatc ttcg | | / |
| | GG5F | ggctacggtctccctttgtagttaagagacacac | 4327 | / |
| | Bsa del orf 1 rev | ggctacggtctcgcctatcagacattatgcaaagtat | | / |
| | BSA del orf 1 fwd | ggctacggtctcgtagggacctttatgacaagttgca | 2866 | / |
| | GG6R | ggctacggtctcgtaatgtgtttaaatattgacacag | | / |

| | | | | |
|---|---|---|---|---|
| | GG7F | ggctacggtctcgattaacattagctgtaccctataatatg | 3282 | / |
| | BsaI spike del rev | ggctacggtctcggtccctagcagcaatatcaccaaggca | | / |
| | BsaI del spike fwd | ggctacggtctcgggacctcatttgtgcacaaaagt | 4392 | / |
| | Orf 8 8Rgg | ggctacggtctcggatgattcctaagaaaacaagaaatttcatgt | | / |
| | Orf8-9Fgg | ggctacggtctcgcatcacaactgtagctgca | 1938 | / |
| | 6R gg | ggctacggtctcgaagctattaaaatcacatggggatagcacta | | / |
| | Link gg fwd | ggctacggtctcggcttcttaggagaatgacaaaaaaaa | 5452 | / |
| | Link rev gg | ggctacggtctcggggaaggtataaacctttaatacggttcactaaaccagctct | | / |
| rN-Delta, rN-BA.2<br>rN-Pirola | CLEVER F1 fwd | cgttacataacttacggtaaatgg | 7747 | / |
| | CLEVER F1 rev | gcagttaaatcccatttaaaagatg | | 101 |
| | CLEVER F2 fwd | ccttgtagtgtttgtcttagtgg | 8090 | |
| | CLEVER F2 rev new | tgttccaattactacagtagctcc | | 160 |
| | CLEVER C fwd | TATAACTCAAATGAATCTTAAGTATGCCATTAGTGCAAAGAATAGAGCTCGCACCGTAGCTGGTG | 5947 | |
| | CLEVER C rev | ATCACCAATCAAAGTTGAATCTGCATCAGAGACAAAGTCATTAAGATCTGAGTCGACAAGCAGCG | | 100 |
| | CLEVER D fwd | TACAGCTGTTTTAAGACAGTGGTTGCCTACGGGTACGCTGCTTGTCGACTCAGATCTTAATGACTTTGTC | 8027 | |
| | CLEVER F4a rev | ctggttactgccagttgaatc | | 84 |
| | N ATG fwd / N ATG delta fwd | atgtctgataatggaccccaaaatca / atgtcttataatggaccccaaaatca | 2162 | |
| | CLEVER F4b rev | tactcaagctttaagatacattgatgagt | | / |

* for N ATG fwd and N ATG Delta fwd - all other primers are the same, but Delta VOC has a mutation at position 3 in the N gene, and so the primer needs to be adapted, other primer constellation stays the same for the rescue

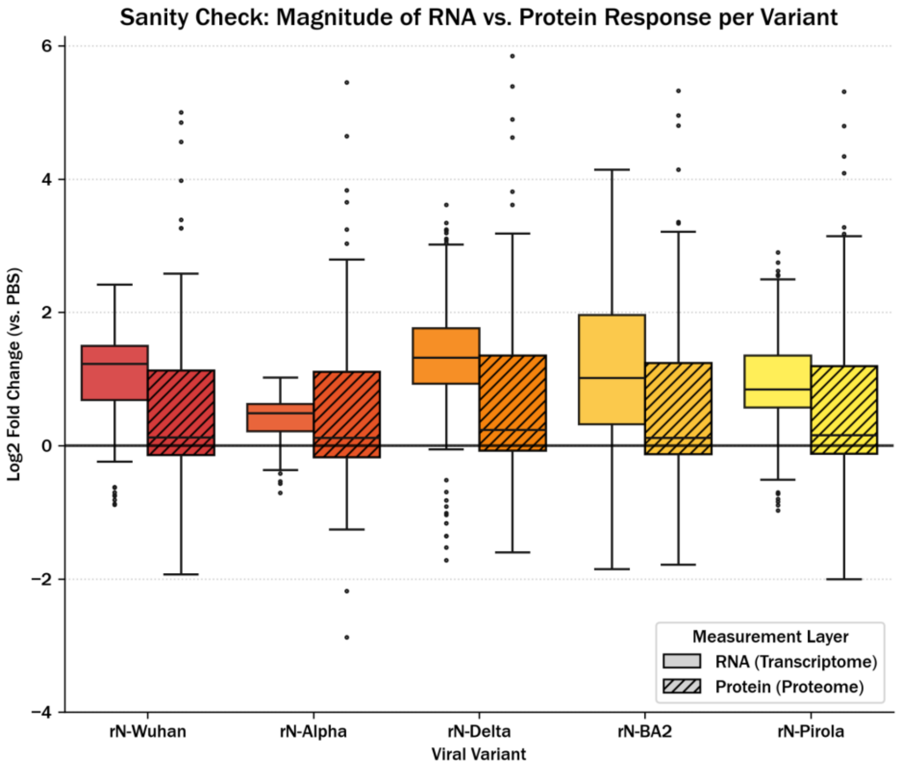

Supplementary Figure 1

**Response magnitude distribution across omics layers.** Grouped boxplot comparing the global distribution of host transcriptomic (RNA, solid fill) and proteomic (protein, hatched pattern) LFC across viral variants relative to the PBS control. Boxplots represent the median and interquartile range, with fliers indicating outliers. The horizontal solid line at zero indicates baseline expression matching the control.

.

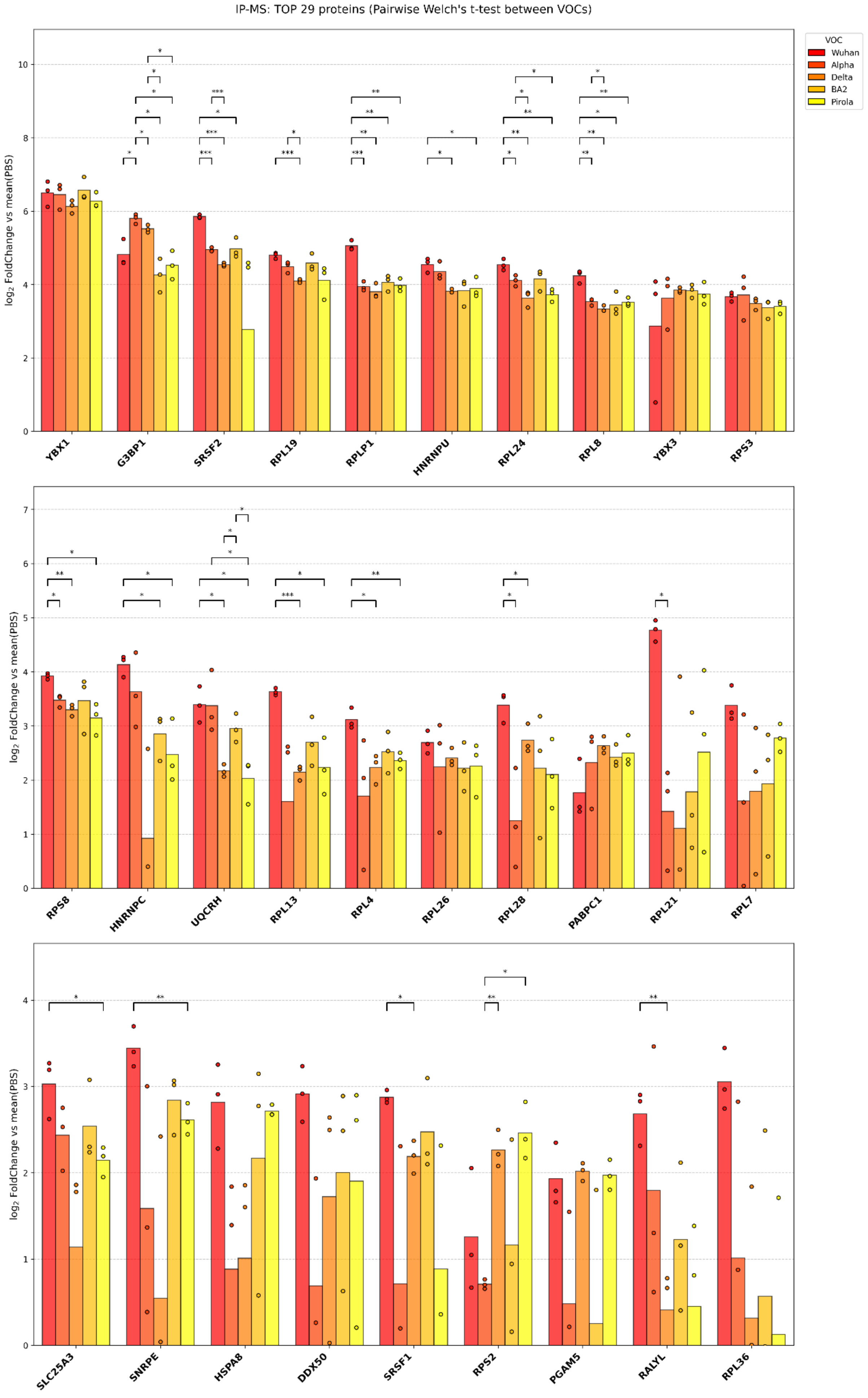
IP-MS: TOP 29 proteins (Pairwise Welch's t-test between VOCs)
VOC
Wuhan
Alpha
Delta
BA2
Pirola
log2 FoldChange vs mean(PBS)
YBX1
G3BP1
SRSF2
RPL19
RPLP1
HNRNPU
RPL24
RPL8
YBX3
RPS3
RPS8
HNRNPC
UQCRH
RPL13
RPL4
RPL26
RPL28
PABPC1
RPL21
RPL7
SLC25A3
SNRPE
HSPA8
DDX50
SRSF1
RPS2
PGAM5
RALYL
RPL36

Supplementary Figure 2

**Differential protein abundance across SARS-CoV-2 rN-VOCs.** Bars indicate the mean LFC of protein abundance for each rN-VOC relative to the mean of the PBS control group. Individual data points represent the single measurements for each replicate (*n* = 3). Only genes exhibiting a significant positive LFC in at least one VOC were included, sorted by their overall mean LFC across variants. Statistical significance between individual VOCs was determined using a pairwise Welch's t-test (two-sided, unequal variances).

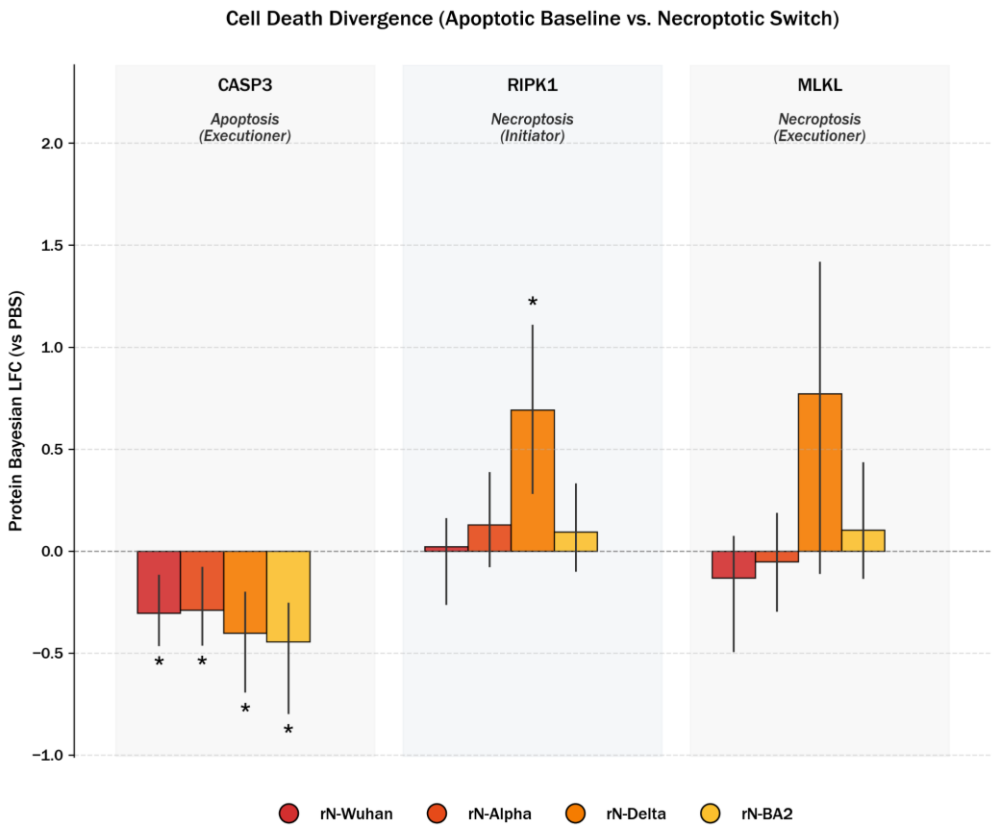

Supplementary Figure 3

A - **Cell death pathway divergence across viral variants in A549-A/T cells.** Error bars indicate the 95% HDI. Asterisks (*) denote statistically significant expression changes where the 95% HDI completely excludes the PBS baseline (normalized to zero).

Supplementary Table 3: Functional clusters IP-MS.

| I: Nucleolus / rRNA Processing | II: Immune Evasion / Stress Granules | III: Splicing / NF-kB Modulation | IV: Mitochondria / Metabolism / Chaperones |
|---|---|---|---|

| | | | |
|---|---|---|---|
| DDX50;RPL4;RPL7;RPL8;RPL13;RPL19;RPL21;RPL24;RPL26;RPL28;RPL36;RPLP1;RPS3;RPS8;SNRPE | G3BP1;PABPC1;YBX1;YBX3;HNRNPU | SRSF1;SRSF2;HNRNPC;RALYL | SLC25A3;'TUFM;UQCRH;PGAM5;HSPA8 |
